# Divergent phenotypes in germline versus conditional mutant mouse models of Sifrim-Hitz-Weiss Syndrome

**DOI:** 10.1101/2023.02.15.528754

**Authors:** Sarah Larrigan, Shrilaxmi Joshi, Pierre Mattar

## Abstract

Chromatin remodellers are among the most important risk genes associated with neurodevelopmental disorders (NDDs), however, their functions during brain development are not fully understood. Here, we focused on Sifrim-Hitz-Weiss Syndrome (SIHIWES) – a brain overgrowth/intellectual disability disorder caused by mutations in the *CHD4* chromodomain helicase gene. We utilized mouse genetics to excise the *Chd4* ATPase/helicase domain – either in the germline, or conditionally in the developing telencephalon. Conditional heterozygotes exhibited little change in cortical size and cellular composition, and had only subtle behavioral phenotypes. Telencephalon-specific conditional knockouts had marked reductions in cortical growth, reduced numbers of upper-layer neurons, and exhibited alterations in anxiety and repetitive behaviors. Despite the fact that germline heterozygotes exhibited comparable growth defects, they were unaffected in these behaviors, but instead exhibited female-specific alterations in learning and memory. These data reveal unexpected phenotypic divergence arising from differences in the spatiotemporal deployment of loss-of-function manipulations, underscoring the importance of context in chromatin remodeller function during neurodevelopment.

**Graphical Abstract.:** 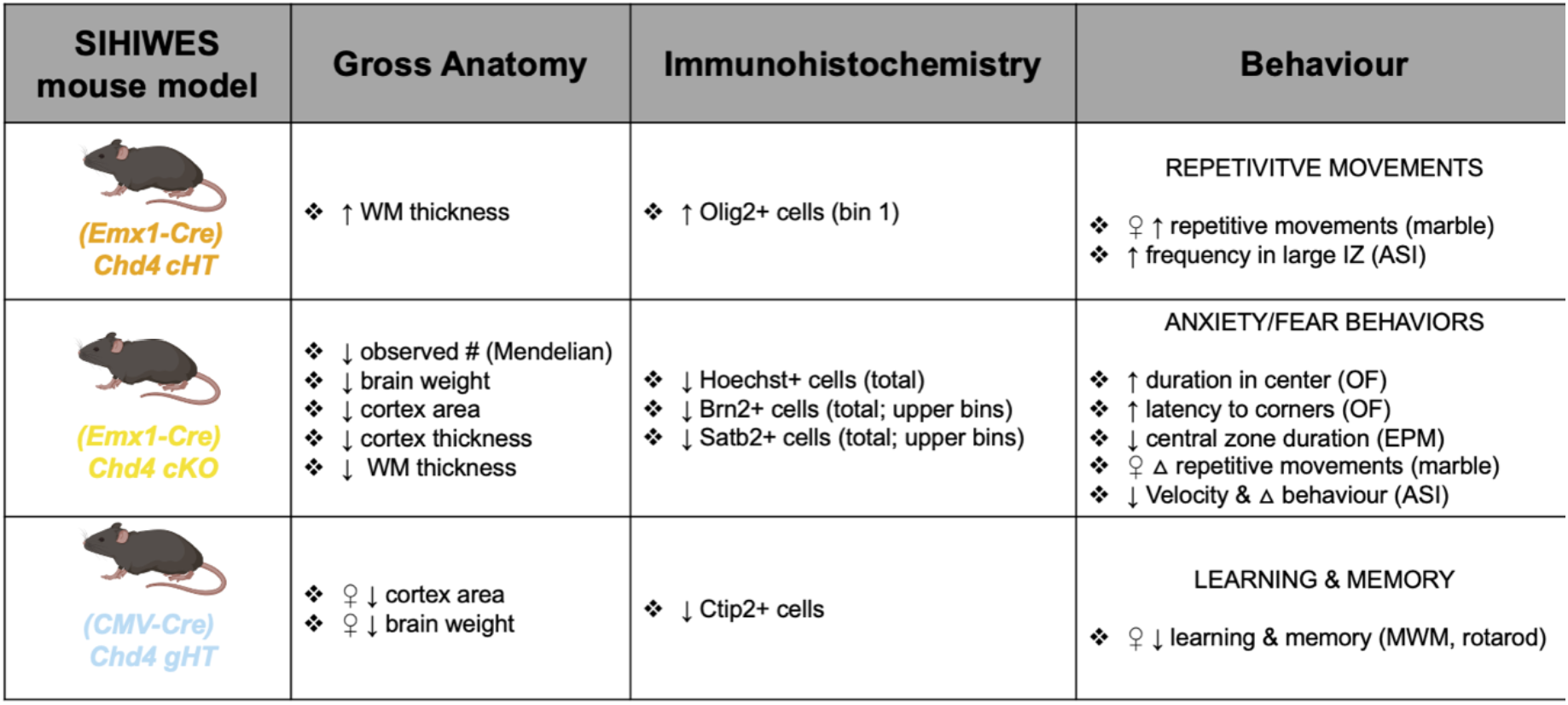

## Introduction

The histogenesis of the central nervous system (CNS) depends upon the ordered deployment of many developmental processes, including progenitor proliferation, differentiation, migration, axon pathfinding, and synapse maturation – all of which critically depend on dynamic changes in gene expression. Chromatin remodelers mobilize nucleosomes to help reconfigure the epigenome, facilitating transitions between activated and repressed states. There are four main families of chromatin remodeling enzymes: SWI/SNF, ISWI, CHD, and INO80, each of which is encoded by multiple paralogues (2). For example, the mammalian CHD family encompasses 10 different genes. Despite this putative redundancy, chromatin remodelers are among the most important risk genes for intellectual disability (ID) and autism spectrum disorder (ASD) (3–5), suggesting that these genes have specific neurodevelopmental functions.

Subfamily II CHD genes – consisting of *CHD3*, *CHD4*, and *CHD5* – are notable. These subunits exclusively incorporate into the nucleosome remodeling and deacetylase (NuRD) complex (6). In addition to nucleosome remodelling activity, the NuRD complex also contains Class I histone deacetylase enzymes (7–9), and thereby functions as an important epigenetic ‘eraser’. CHD4 has been additionally shown to incorporate into a second complex called ChAHP (CHD4, ADNP, HP1) (10), which does not include histone deacetylases. Both NuRD and ChAHP complexes are associated with gene repression, heterochromatinization, and suppression of genomic accessibility (6, 10–15).

*De novo* variants in *CHD4* cause the overgrowth-intellectual disability syndrome known as Sifrim-Hitz-Weiss syndrome (SIHIWES; MIM 617159) (16–18). 32 individuals with SIHIWES have been characterized to-date. All patients were heterozygous with nearly all variants being missense or in-frame insertions/deletions, with the exception of three variants predicted to lead to truncated proteins. Notably, the majority of the missense mutations mapped to the ATPase/helicase enzymatic domains. SIHIWES symptoms are variable in affected patients. Mild to moderate ID, macrocephaly, and delayed speech are hallmark symptoms. Additionally, 22 of the 23 individuals that underwent MRI brain imaging presented with neuroanatomical abnormalities including ventriculomegaly, hydrocephalus, Chiari 1, and corpus callosum hypoplasia. Outside of the brain, SIHIWES was also associated with hearing loss, hypotonia, motor delay, and abnormalities in the heart, skeleton, vasculature, and gonads (16–18). These findings support the idea that CHD4 acts on basic developmental processes. Other subfamily II genes have also been linked to neurodevelopmental disorders, with variants in the *CHD5* and *CHD3* genes identified as the cause of Parenti-Mignot neurodevelopmental syndrome (MIM 619873) and Snijders Blok-Campeau syndrome (MIM 618205), respectively (19, 20).

To begin to understand the molecular mechanisms through which CHD4 contributes to neurodevelopment, several animal models have been generated. *Chd4* is an essential gene (1, 21), necessitating conditional approaches to generate CNS knockouts. In postmitotic cerebellar neurons, conditional knockouts (cKOs) have revealed that *Chd4* plays critical roles in gene regulation and circuit function (14, 22, 23). With respect to growth of the brain, previous work has shown that *Chd4* is required for the survival and proliferation of cortical progenitor cells during neurogenesis (24), but perinatal lethality precluded analysis of postnatal brain development and behavior. Moreover, while *CHD4* mutations associated with SIHIWES are heterozygous, the reported phenotypes in both the cerebellum and cortex were observed only in homozygous *Chd4* cKOs. Heterozygote phenotypes were not reported in the above studies.

To compare how *Chd4* knockout and/or haploinsufficiency affect neurodevelopment, we generated novel mouse models. Using the *Emx1-Cre* driver (25), we conditionally ablated *Chd4* from the developing dorsal telencephalon. Conditional heterozygotes (cHTs) and cKOs proved to be viable, permitting us to examine postnatal phenotypes. We additionally used *CMV-Cre* (26) to create germline heterozygotes (gHTs). While we expected *Chd4* cHT and gHT mice to be similar, cHT mice exhibited behavioral phenotypes related to sociability and repetitive behaviors. By contrast gHT mice had significant reductions in cortical size, and divergent behavioral phenotypes. Our data reveal surprising phenotypic heterogeneity arising from similar loss-of-function manipulations, underscoring the pleiotropy of chromatin remodellers in neurodevelopment.

## Results

In order to assess the role of *Chd4* in neurodevelopment and behavior, we began by examining Chd4 expression. In accordance with Nitarska et al. (24), we found that Chd4 protein was expressed in a widespread fashion throughout cortical development – both in dividing progenitor cells, and in differentiated neurons and glia (Fig. S1).

As most patients with SIHIWES harbor mutations in the enzymatic domains of *CHD4* (17), we took advantage of a previously developed mouse line that allows for excision of the *Chd4* ATPase/helicase domain using a Cre/loxp approach (1). We utilized *Emx1-Cre* (25) to ablate *Chd4* in the developing dorsal telencephalon, generating cHTs and cKOs. We additionally used *CMV-Cre* (26) to ablate *Chd4* in the germline, generating gHTs (Fig. 1A). To validate the conditional genetics strategy as well as our Chd4 immunohistochemistry, we first examined Chd4 protein expression using 3 antibodies, 2 of which were raised against epitopes mapping outside of the floxed ATPase/helicase domain. Staining revealed that Chd4 protein was lost in the embryonic *Chd4* cKO neocortex as expected (Fig. 1B, C, see also Fig. 3).

**Figure 1.**
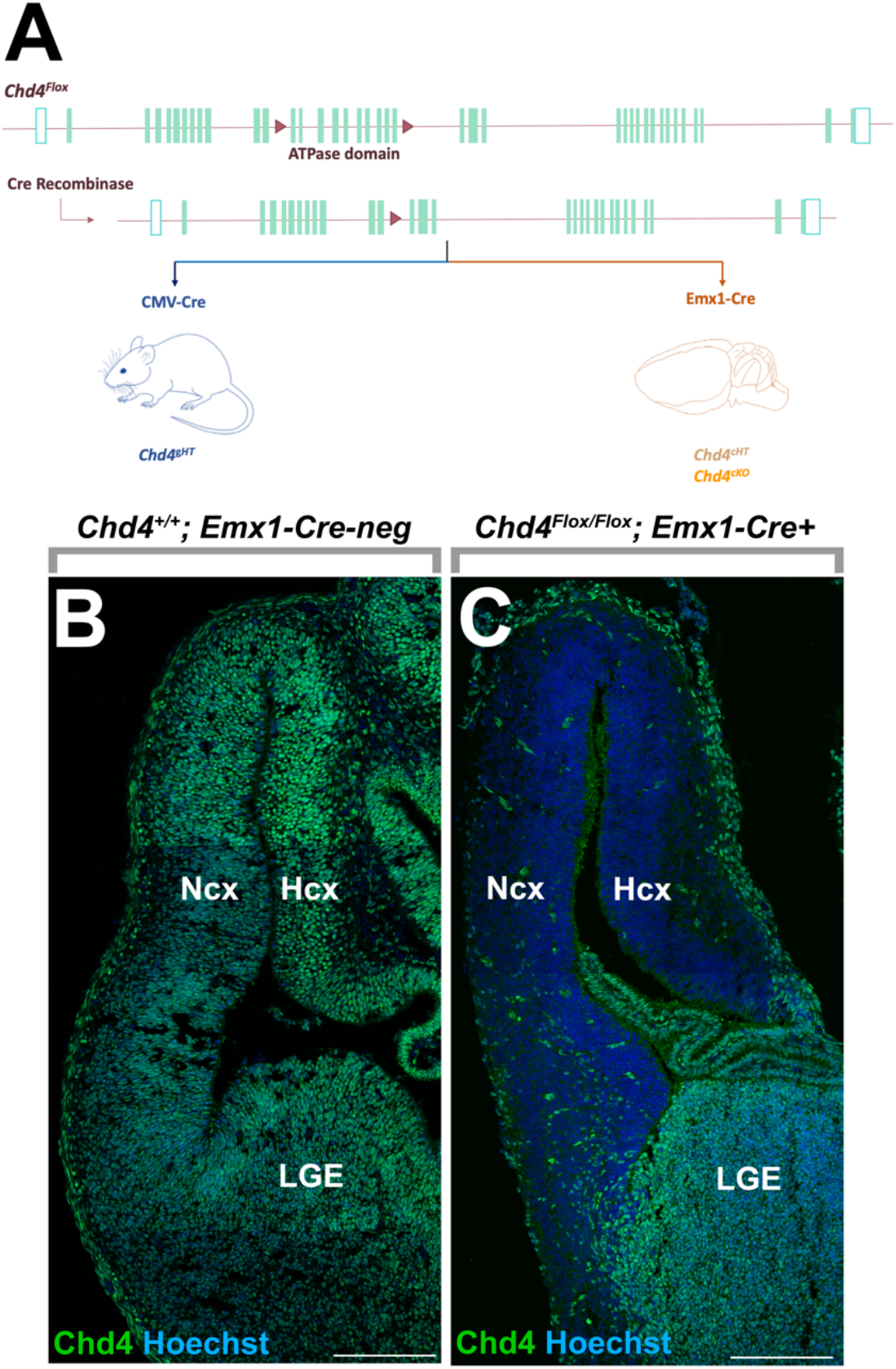
*Chd4* conditional and germline mutants. (A) Genetic strategies for generating telencephalon-specific *Chd4^flox/+^* conditional heterozygotes (cHTs) and *Chd4^Flox/Flox^* conditional mutants (cKOs), as well as germline heterozygotes (gHTs). The *Chd4^Flox^* allele has loxP sites flanking exons 12 to 21, encompassing the ATPase domain (1). (B, C) Chd4 immunohistochemistry on E13.5 cHT (B) and cKO (C) forebrains. Scale bar = 200 μm.

### Chd4 is required for cortical growth

*Chd4* cKOs had previously been generated using the *Nestin-Cre* driver allele, but were reported to be perinatally lethal (24). We reasoned that the later onset and more spatially restricted expression of the *Emx1-Cre* driver might improve the survival of cKOs. Indeed, we were able to successfully generate adult *Chd4* cKOs. However, we found that they were not recovered in Mendelian frequencies suggesting considerable developmental lethality (Table 1). *Chd4* gHT mice were born in normal Mendelian ratios.

**Table 1.** Postnatal survival of *Chd4* cHT and cKO mice. *Emx1-Cre+; Chd4^Flox/+^* mice were crossed with *Chd4^Flox/Flox^* mice to yield *Chd4* cHTs and cKOs. Litters were collected between P7 and P21 (77 pups total). Probability based off Chi-square value was calculated. Degrees of Freedom = 3.

| *Emx1-Cre*<sup>+</sup>; *Chd4*<sup>Flox/+</sup> x *Chd4*<sup>Flox/Flox</sup> | *Chd4*<sup>Flox/Flox</sup> |  | *Chd4*<sup>Flox/+</sup> |  |
| --- | --- | --- | --- | --- |
|  | *Emx1-Cre*<sup>-</sup> | *Emx1-Cre*<sup>+</sup> | *Emx1-Cre*<sup>-</sup> | *Emx1-Cre*<sup>+</sup> |
| **Expected** | 19.25 | 19.25 | 19.25 | 19.25 |
| **Observed** | 34 | 3 | 14 | 26 |
| **Chi-square value** | 11.30195 | 13.71753 | 1.431818 | 2.366883 |
| **Probability** | 0.05* | 0.01** | n.s. | n.s. |

Next, we measured brain sizes and weights at postnatal day (P) 7. Grossly, we found that *Chd4* cKOs had a marked reduction in the size of the dorsal telencephalon (Fig. 2A). Brain weight was accordingly reduced in cKO animals (Fig. 2B). *Chd4* cHT brains were not significantly different versus controls, however, *Chd4* gHTs had a significant reduction in brain weight (Fig. 2B). Similar differences were observed in cortical area - measured by manually tracing the telencephalic hemispheres in two-dimensional images (Fig. 2C). To determine whether these growth defects were specific to the cortex, we performed similar area measurements of the cerebellum. We found no difference in cHT and cKO cerebellar area, in accordance with the lack of *Emx1-Cre* expression in this territory (25). The gHT cerebellum area was likewise unaffected despite body-wide heterozygosity (Fig. 2D). As an additional control, we measured animal body weights. We found no significant difference between control, cHT, and cKO animals, although weights of cKO animals were significantly reduced in comparison to cHTs. Likewise, gHT animals were not different from wild-type controls (Fig. 2E). Thus, we find that *Chd4* is required for cortical growth, as was previously described by the Riccio laboratory (24). Interestingly, we find that in the hemizygous condition, *Chd4* can also lead to cortical hypoplasia when deleted in the germline, but not when deleted conditionally at the onset of neurogenesis.

**Figure 2.**
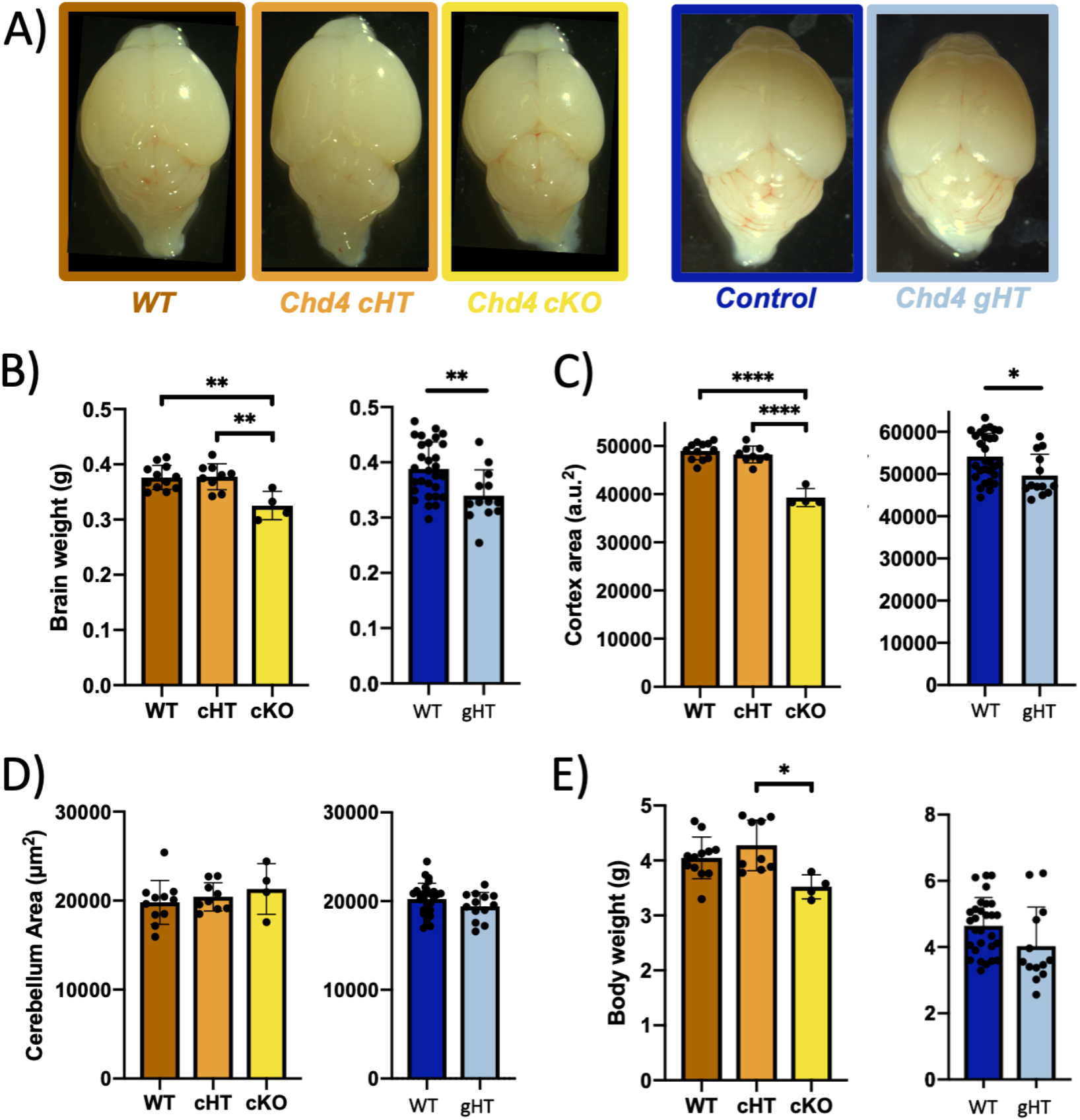
*Chd4* is required for cortical growth. (A) Wholemount images of wild-type, cHT, cKO, and gHT brains harvested at P7. (B-E) Size measurements according to genotype as indicated. (B) Brain weight. (C) Cortex area. au: arbitrary units. (D) Cerebellum area. (E). Body weight. Statistical analyses for WT, cHT, and cKO comparisons are via one-way ANOVA with Tukey’s post-hoc test. Comparisons of control vs. gHT are via Student’s t test. ^*^ p < 0.05; ^**^ p <0.01; ^****^ p <0.0001. *Emx1-Cre* line: WT n=11, cHT n=9, cKO n=4. *CMV-Cre* line: WT n=29, gHT n= 13.

To determine how the gross changes in cortical size relate to cellular composition, we next examined P7 brains histologically, focusing on the somatosensory cortex. We first performed immunohistochemistry for Chd4 in coronal tissue sections. As per embryonic stages (Fig. 1C), Chd4 protein expression was markedly reduced in the *Chd4* cKO (Fig. 3A). Although residual Chd4 protein remained expressed, the observed pattern was consistent with expression from inhibitory interneurons and endothelial cells that are outside the *Emx1-Cre* lineage. Chd4 protein expression in cHT and gHT brains was not obviously different versus controls (Fig. 3A).

**Figure 3.**
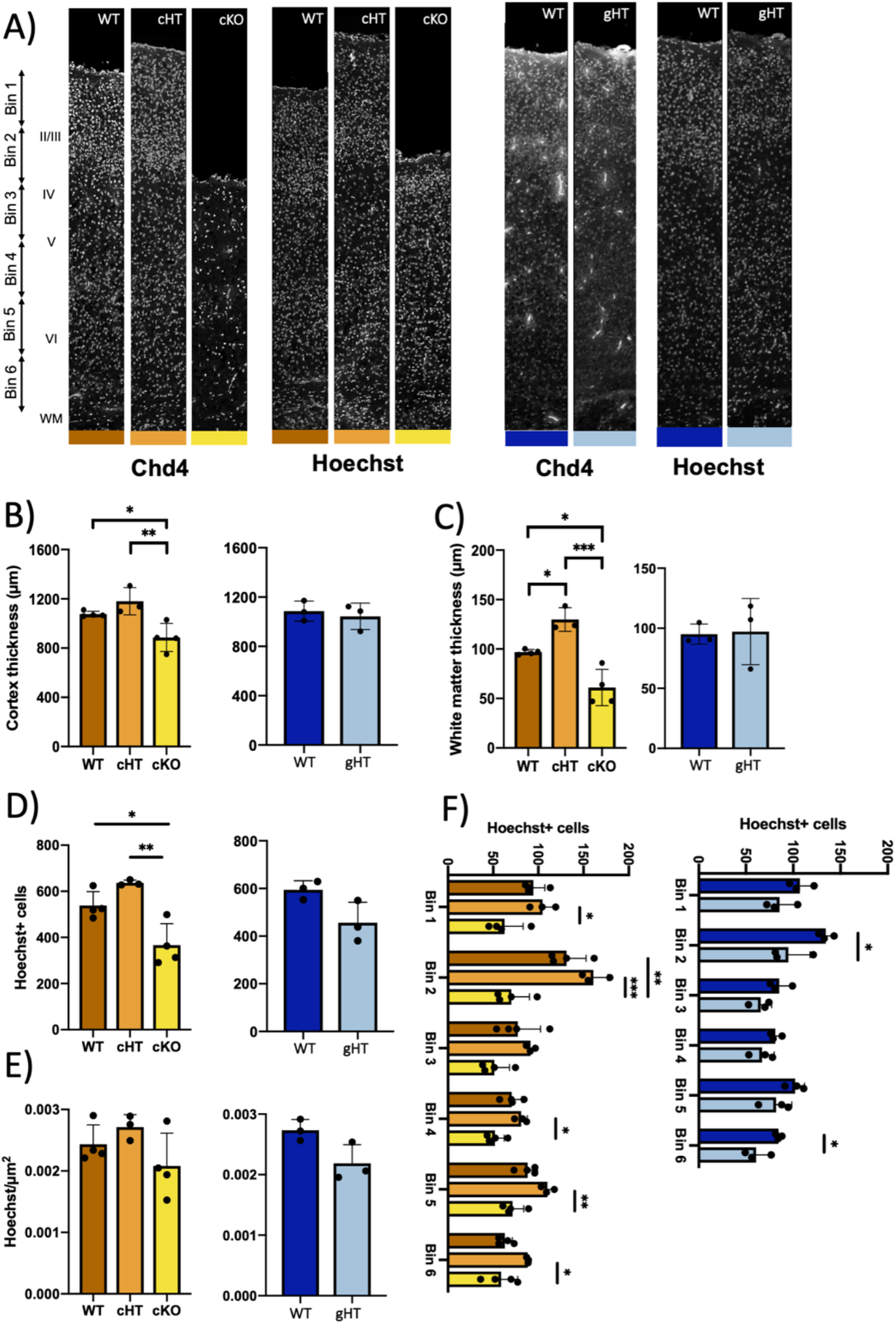
Changes in cell number in *Chd4* mutants. (A) Immunohistochemistry on coronal sections from wild-type, cHTs, cKOs, or gHTs for Chd4 protein, harvested at P7. (B, C) Measurements of the cross-sectional thickness of the cortical plate (B) or white matter (C). (D-G) Cell counts within the cortical plate according to genotype as indicated. (D) Absolute count of Hoechst+ cells. (E) Cell density. (F) Absolute counts of Hoechst+ cells taken from 6 equally sized bins (see A). Statistical analyses for WT, cHT, and cKO comparisons are via one-way ANOVA with Tukey’s post-hoc test. Comparisons of control vs. gHT are via Student’s t test. ^*^ p < 0.05; ^**^ p <0.01; ^***^ p <0.001. *Emx1-Cre* line: WT n=4, cHT n=3, cKO n=4. *CMV-Cre* line: WT n=3, gHT n=3.

We next measured the thickness of the neocortical layers. In accordance with the gross reduction in cortical size observed in *Chd4* cKOs, we found that the thickness of the neuronal layers was significantly reduced (Fig. 3B). While cHT neocortices appeared to be thicker versus controls (Fig. 3A), neuronal layers were not significantly differently (Fig. 3B). Instead, this effect appeared to be due to an increase in the thickness of the white matter (inner fiber layer), which was significantly expanded in cHTs relative to controls (Fig. 3C). Conversely, white matter thickness was significantly reduced in cKOs (Fig. 3C). gHT mice were not different from controls.

These observations were echoed in cell counts. We quantitated Hoechst+ nuclei in 200μm-wide columns of the somatosensory cortex, which revealed decreases in cKOs (Fig. 3D), and no change in gHT cell numbers. Moreover, there were no changes in cell density in any of the genotypes (Fig. 3E). Next, dividing the neuronal layers into 6 equally sized bins, we found that the reduced cell number exhibited by *Chd4* cKOs was most prominent within bin 2 (Fig. 3F), which generally maps to the upper layers of the cortex (see Fig. 3A). Cell counts in gHT neocortices were significantly reduced in 2 out of 6 bins (Fig. 3F), perhaps reflecting the decrease in gross cortical size observed previously in wholemount (Fig. 2A, B).

### Reductions in upper-layer neurons in Chd4 mutants

Next, we examined cell-type specific marker proteins. We examined markers for lower layer (Ctip2 and Tbr1) and upper-layer (Brn2 and Satb2) neuronal subtypes (Fig. 4A). We again counted cells within 200μm-wide columns, dividing the neuronal layers into 6 equally-sized bins as per above. In accordance with the reduction in total cell number, we found a significant reduction in upper-layer neurons in the cKO – marked both by Brn2 and Satb2 (Fig. 4B, C).

**Figure 4.**
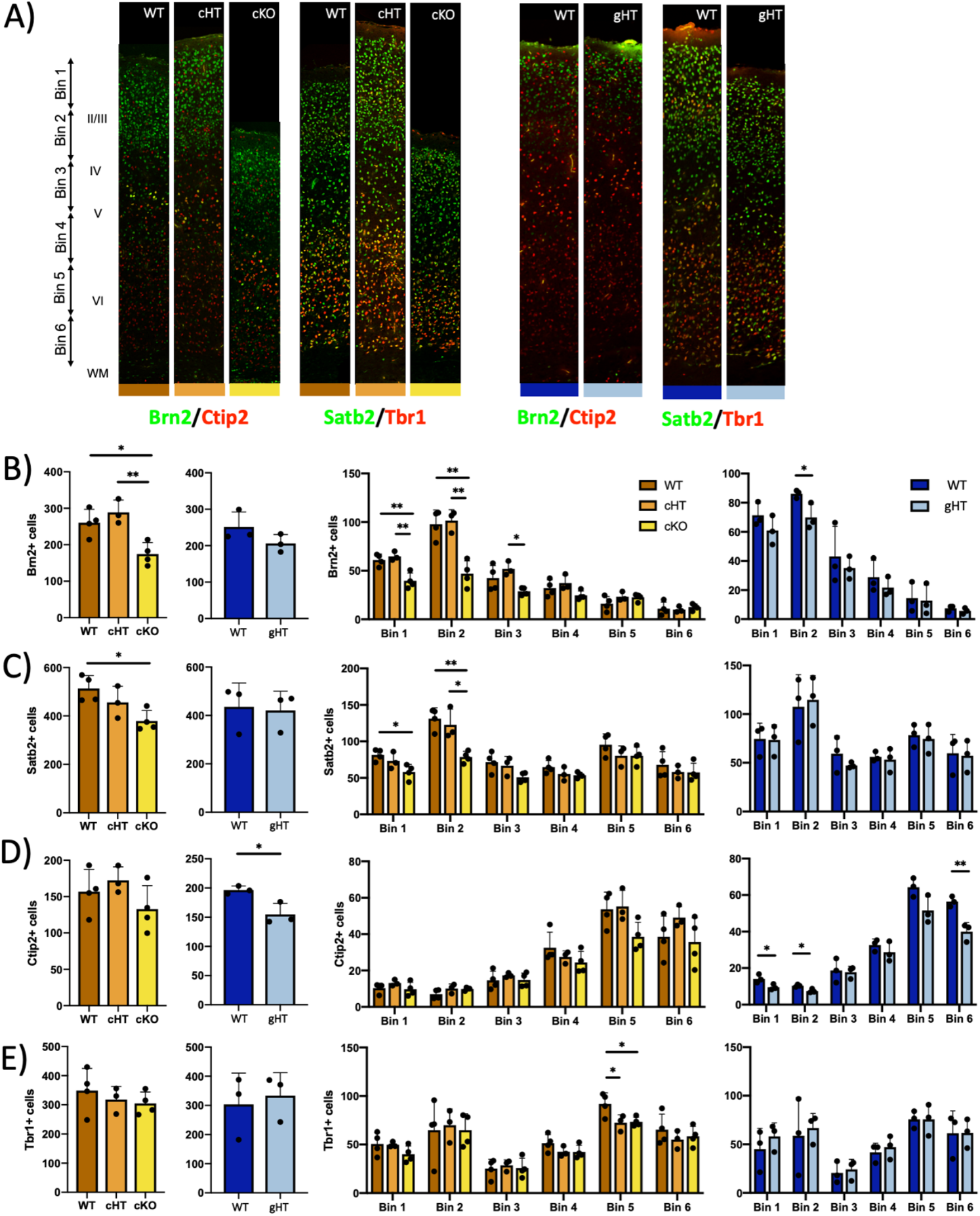
*Chd4* is required to expand upper neocortical layers. (A) Immunohistochemistry on wild-type, cHTs, cKOs, or gHTs for the upper-layer markers Brn2 and Satb2, or the lower-layer neuron markers Ctip2 and Tbr1. (B-E) Absolute cell counts, or counts taken from 6 equally sized bins (see A) for the indicated genotypes. (B) Brn2. (C) Satb2. (D) Ctip2. (E) Tbr1. Statistical analyses for WT, cHT, and cKO comparisons are via one-way ANOVA with Tukey’s post-hoc test. Comparisons of control vs. gHT are via Student’s t test. ^*^ p < 0.05. ^**^ p < 0.01. *Emx1-Cre* line: WT n=4, cHT n=3, cKO n=4. *CMV-Cre* line: WT n=3; gHT n=3.

We found that early born lower-layer cortical neurons, marked by Ctip2 and Tbr1, were not affected in cHTs and cKOs (Fig. 4D, E). However, we observed a significant reduction in Ctip2+ neurons in gHT neocortices (Fig. 4D). Ctip2 expression is most prominent in Layer V, but we did not observe a difference in Ctip2 expression in the corresponding bin (bin 5). Instead, reductions were most apparent in the most basal bin 6, as well as in the most apical bins 1 and 2 (Fig. 4D), where Ctip2 is normally expressed at low levels. Moreover, these changes were not matched by changes in Tbr1+ cell numbers (Fig. 4E). These data suggest that the observed changes in Ctip2 expression may not be driven by underlying changes in neuron numbers, but might instead reflect alterations in Ctip2 expression levels.

Finally, we measured glial cell numbers, but observed no differences in overall numbers of Olig2+ oligodendrocyte-lineage cells, or in Aldh1l1+ astrocytes (Fig. S2). However, we found that cHTs had significant increases in Olig2+ cells in specific cortical bins, perhaps in accordance with the white matter expansion observed in these mutants. Together, these data indicate that *Chd4* is specifically required for the expansion of late-born cortical neurons, and suggest that *Chd4* germline haploinsufficiency may additionally affect Ctip2 expression.

### Assessing activity and anxiety levels in Chd4 mutants

To determine how the observed anatomical changes correlated with behavioral defects, we ran mutant mice through a battery of tests. We chose to assess *CMV-Cre* and *Emx1-Cre* lines independently due to subtle but significant differences observed in the baseline behavior of the control mice. We examined sexes separately, but generally found few differences and therefore pooled male and female data together in order to increase statistical power. However, we present sex-separated data where significant differences were observed.

As many of our behavioral tests depend on vision, we tested the optokinetic reflex of mutant mice, but observed no differences versus controls (Fig. S3).We measured circadian activity levels using the beam-break test, in which mice are placed in a cage containing an array of photobeams. The ambulatory movement of mice within the cage leads to beam interruptions that are tabulated over the course of 48 h. We observed no difference in the circadian activity levels of *Chd4* cHT, cKO, or gHT mice versus controls (Fig. 5A, B).

**Figure 5.**
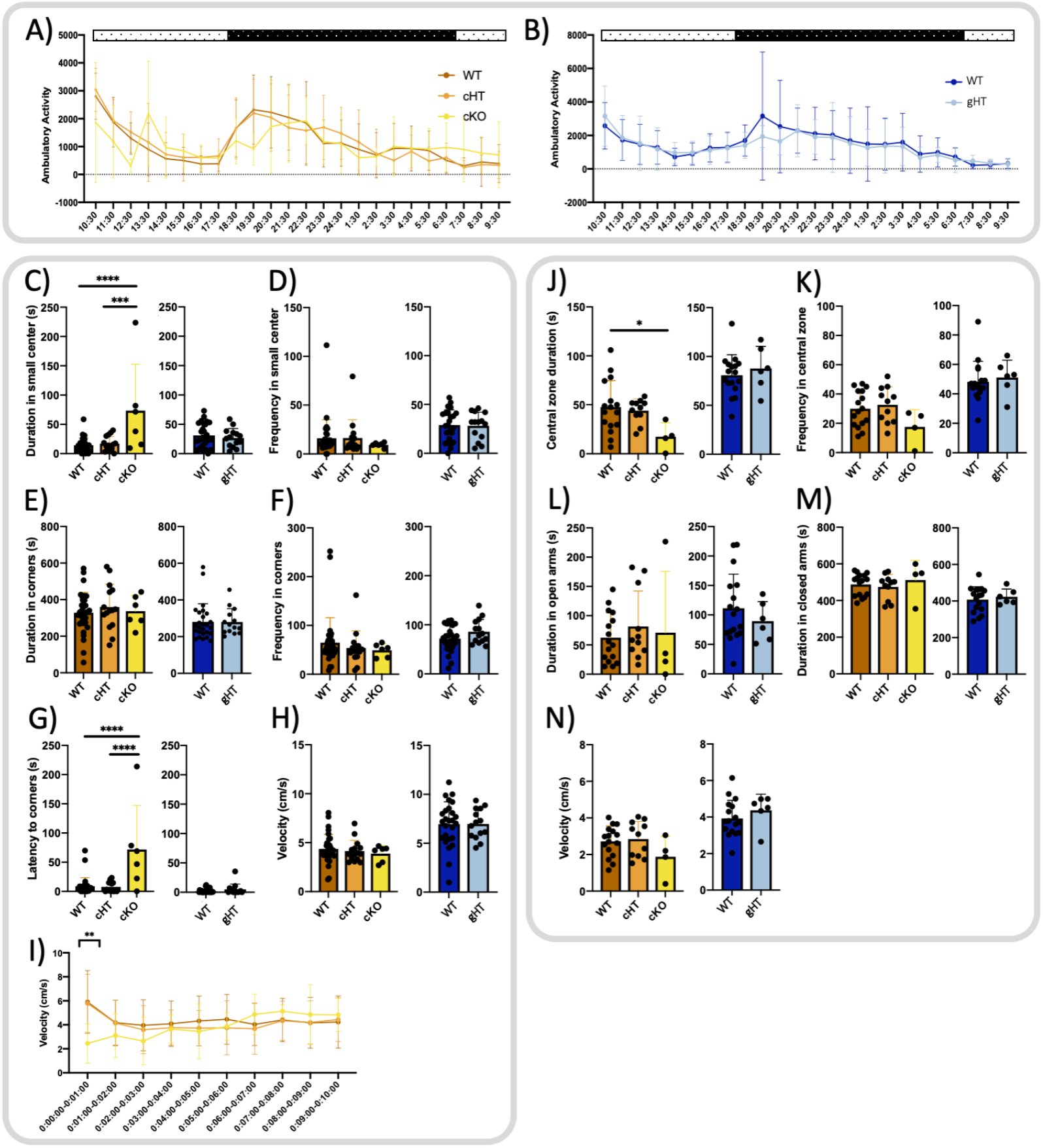
Exploration, activity levels, and anxiety behaviors in *Chd4* mutants. (A, B) Cumulative ambulatory activity counts per hour in the beam break assay for (A) wild-type (n=32), cHT (n=15), and cKOs (n=6), or (B) wild-type (n=26) and gHTs (n=16). Stippled bars indicate the light/dark cycle. (C-I) Open field test for wild-type (n=32), cHT (n=15), and cKOs (n=6), or wild-type (n=26) and gHTs (n=14) as indicated. Cumulative duration (C) and frequency (D) in the small center. Cumulative duration (E) and frequency (F) in the corners. Latency to reach the corners (G). Average velocity throughout the entire testing interval (H) or broken down into 1-minute intervals (I). (J-N) Elevated plus maze test for wild-type (n=16), cHT (n=11), and cKOs (n=4), or wild-type (n=17) and gHTs (n=13) as indicated. Cumulative duration (J) and frequency (K) in the central zone. Cumulative duration in the open arms (L) or closed arms (M). (N) Average velocity throughout the testing interval. Statistical analyses for WT, cHT, and cKO comparisons are via one-way ANOVA with Tukey’s post-hoc test. Comparisons of control vs. gHT are via Student’s t test. ^*^ p < 0.05; ^**^ p <0.01; ^***^ p <0.001; ^****^ p < 0.0001.

Having found no differences in gross sensorimotor behavior, we examined mutant mice in the open field test, which assesses activity levels and anxiety. Animals are placed in a square-shaped arena with brighter illumination at the center. Mice are normally light-aversive, and spend more time in the darker corners of the chamber. *Chd4* cHT and gHT mice did not exhibit differences versus controls. However, we observed that *Chd4* cKOs spent more time in the open central quadrant of the testing arena (Fig. 5C). Upon further examination, we determined that the *Chd4* cKO phenotype arises due to the fact that the mice ‘freeze’ upon being placed in the testing arena, leading to increased latency to reach the corners. While overall velocity levels were not affected (Fig. 5H), *Chd4* cKOs exhibited reduced velocity specifically within the first minute of testing (Fig. 5I), after which, their activity patterns were indistinguishable from controls. These data suggest that cKO mice may experience elevated anxiety upon encountering a novel environment.

We observed similar phenomena in the elevated plus maze test – which also tests for fear/anxiety. Mice are placed in a ‘plus symbol’ -shaped maze containing two walled arms and two open arms. Mice typically prefer the more sheltered closed arms of the maze, and avoid the open arms and the central zone. In these respects, *Chd4* cHT and gHT mice did not differ from controls (Fig. 5J-M). However, we found that *Chd4* cKOs spent significantly less time in the central zone (Fig. 5J). While these observations might suggest that *Chd4* cKOs are less active than controls, we did not observe a significant decrease in velocity (Fig. 5N), ruling out this possibility. Taken together, these results suggest that *Chd4* cKOs may have elevated anxiety levels in novel and/or open environments.

### Assessing social and repetitive behaviors in Chd4 mutants

We performed behavioral testing for social and repetitive behaviors, which are often affected in mouse NDD models. In the adult social interaction test, a mouse is first habituated to an empty cage containing an inner chamber. Test mice are then reintroduced into the cage with a ‘stranger’ mouse present in the inner chamber. *Chd4* cKOs exhibited marked differences in this test, although they likely do not reflect a phenotype in sociability. When stranger mice were placed in the interaction chamber, we found that *Chd4* cHT, cKO, and gHT mice spent more time in the interaction zone of the testing arena as expected (Fig. 6A, B). However, *Chd4* cKOs entered and occupied the socialization areas significantly less frequently. Importantly, the same effect occurred during the habituation phase, when the interaction partner mouse was absent (Fig. 6A, B). We found the ratio of time spent in the interaction zone in the habituation versus socialization phases was not significantly different versus controls (Fig. 6C). Instead, we found that *Chd4* cKOs exhibited decreased velocity throughout both the habituation and socialization phases of this test (Fig. 6D, E). This suggests that the environment of this particular test affected the cKO mice such that they were more hesitant to explore the testing arena – again suggesting an anxiety-related phenotype.

**Figure 6.**
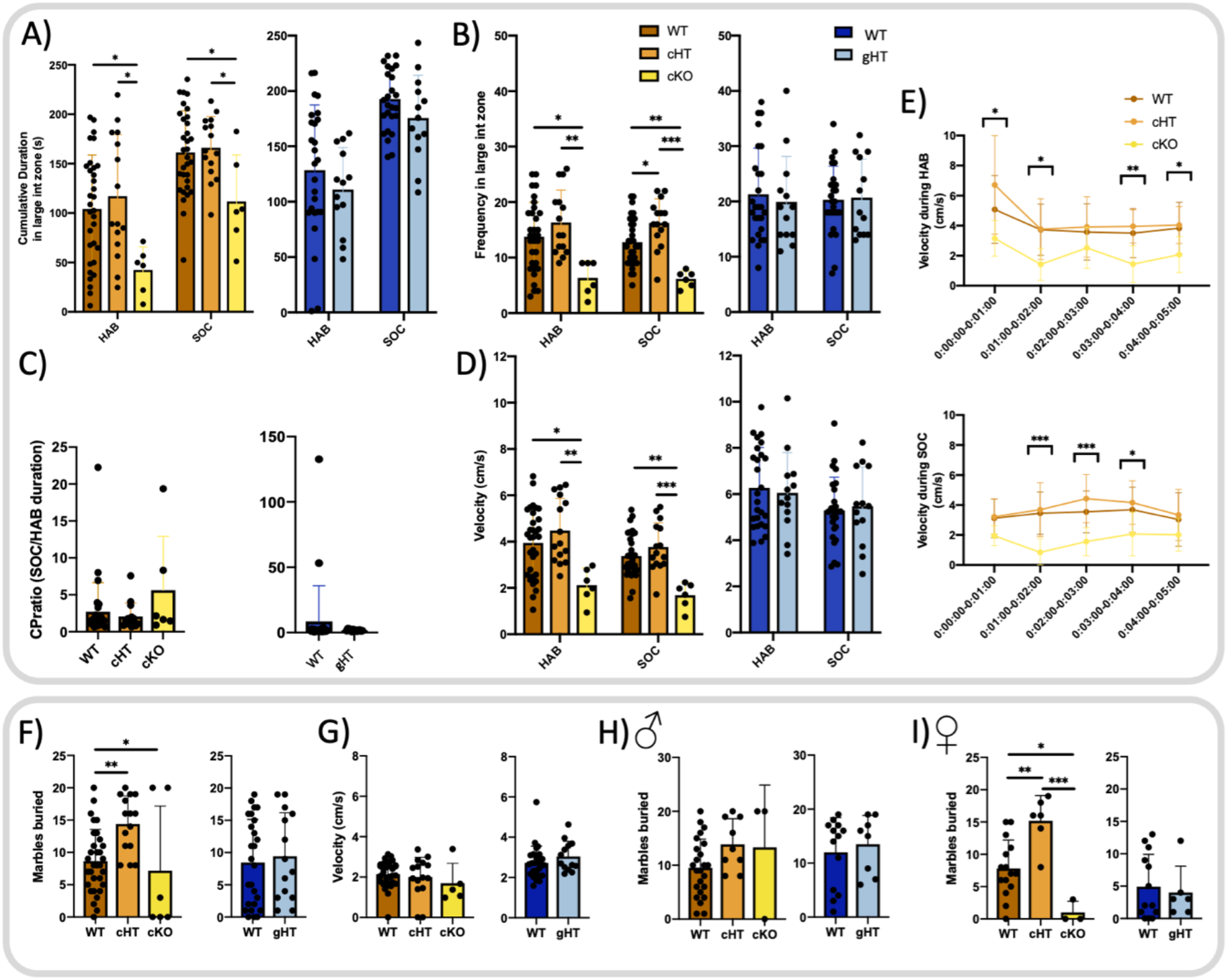
Sociability and repetitive behaviors in *Chd4* mutants. (A-D) Adult social interaction test, comparing the habituation phase (when the partner mice are absent) versus the socialization phase (when interaction partner mice are introduced into the chamber). Cumulative zonal duration (A), frequency (B), ratios of zonal duration (C), distance traveled (D), and velocity (E) for wild-type (n=32), cHT (n=15), and cKOs (n=6), or wild-type (n=26) and gHTs (n=13). (F-I) Marbles buried (F) and velocity (G) during the marble burying test for wild-type (n=32), cHT (n=15), and cKOs (n=6), or wild-type (n=26) and gHTs (n=14). (H, I) Shows the marble burying data shown in (F, G), but separated by sex. Wild-type (n= 13♂ and 13♀) or gHT (n= 8♂ and 5♀). Statistical analyses for A-C, E, and F are via One-way ANOVA with Tukey’s post-hoc test. Statistical analyses for D and G are via unpaired Student’s t-test. ^*^ p < 0.05; ^**^ p <0.01.

While *Chd4* cHTs spent equivalent time in the interaction zones versus controls (Fig. 6A), we found that they entered the large interaction zone around the stranger mouse significantly more frequently during the socialization period compared to wildtype, and not during the habituation phase (Fig. 6B). From this, we deduced that *Chd4* cHTs visit the stranger mouse more frequently but stay for briefer periods of time, which we later confirmed with video footage, suggesting a relatively subtle but nonetheless statistically significant alteration in social behavior.

To examine stereotyped and repetitive behaviors, we performed the marble burying test. Mice are introduced into a cage containing 20 marbles, and the number of marbles buried in a 30-minute period is recorded. We found that *Chd4* cHTs buried significantly more marbles compared to controls (Fig. 6F), suggesting increased repetitive behaviors. By contrast, *Chd4* cKOs buried nearly all the marbles, or none at all. Overall, a statistically significant decrease was observed despite high variance (Fig. 6F). Interestingly, for both cHT and cKO animals, we found that the phenotype appeared to be driven by the performance of female but not male animals (Fig. 6H, I). Taken together, the adult social interaction and marble burying tests reinforce the notion that *Chd4* cKOs exhibit elevated anxiety in specific environments, and implicate both cHTs and cKOs in repetitive behaviors. Despite the fact that gHTs would be expected to capture all of the phenotypes exhibited by cHTs, we observed no social or repetitive behavioral phenotypes in gHTs.

### Germline Chd4 heterozygotes exhibit sex-specific learning deficits

To examine learning and memory, we performed the Morris water maze test (Fig. 7A). Mice were trained over the course of 6 days to find a platform based on visual cues. On the 7th day, (probe day) the platform was removed, which tests the ability of the mice to remember the platform’s location. Neither *Chd4* cHTs or cKOs presented with significant changes in learning and memory during the test (Fig. 7B), although *Chd4* cKOs were significantly different versus cHTs in crossing the platform area on probe day (Fig. 7B). When examined separately, males and females behaved equivalently (Fig. 7C, D). Fascinatingly, whereas *Chd4* gHT mice did not demonstrate significant changes in activity, anxiety, repetitive, or social behaviors (Figs. 5, 6), they displayed significant deficits in learning and memory. From day 3 of the training phase onward, *Chd4* gHT mice took longer to find the platform than wild-type control mice (Fig. 7E). This was particularly evident in female gHT mice (Fig. 7F, G). Additionally, female gHTs had significantly decreased frequency of crossing the platform area on probe day (Fig. 7G).

**Figure 7.**
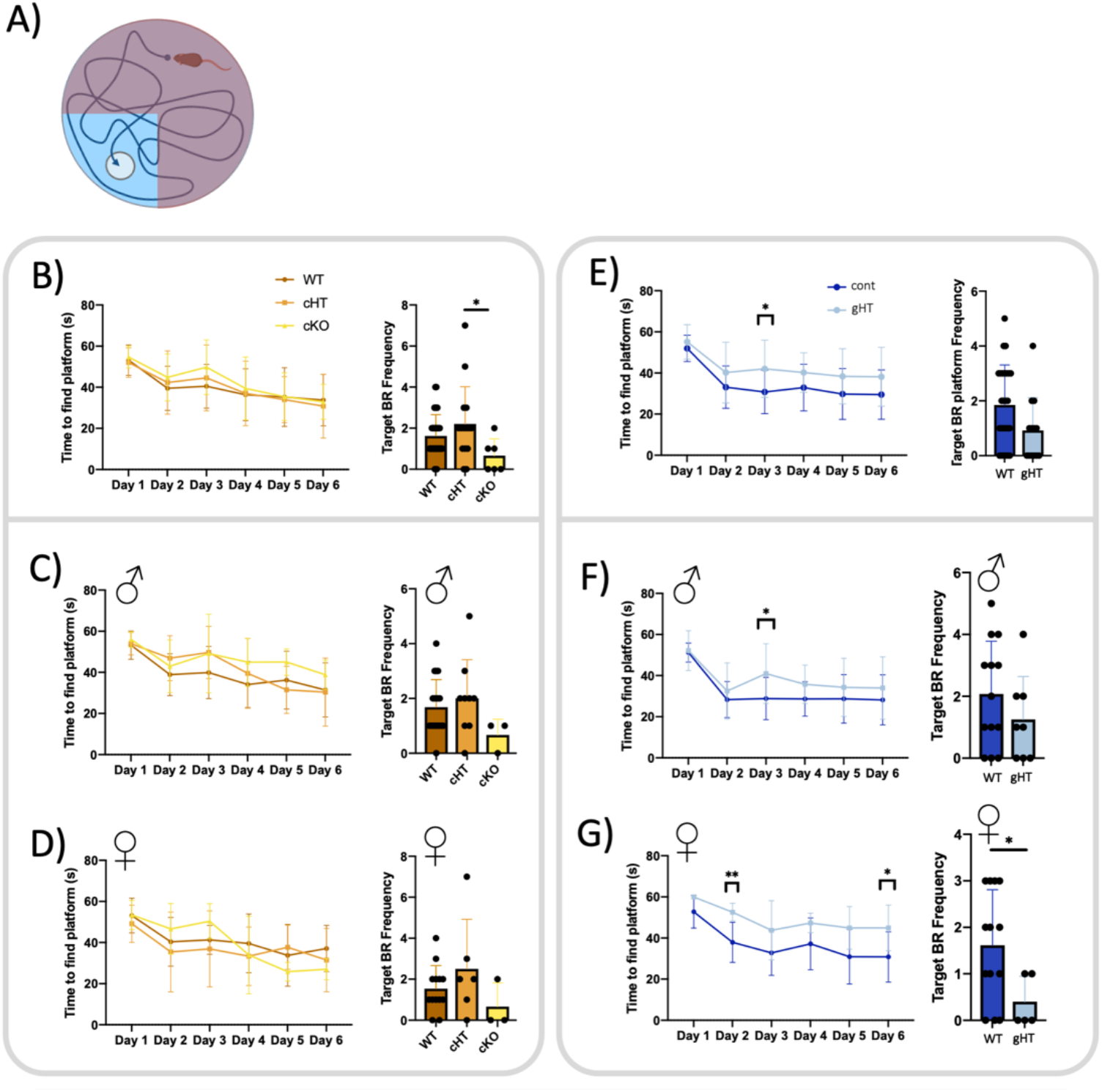
Sex-specific learning, memory, and growth deficits in *Chd4* gHTs. (A) Schematic of Morris water maze test. (B-G) Latency to find the platform during training, and frequency to find the hidden platform on probe day in the Morris water maze test. (C, D, F, G) show the data shown in (B, E), but separated by sex. *Emx1-cre line*: wild-type (n= 19♂ and 13♀), cHT (n= 9♂ and 6♀), and cKO (n= 3♂ and 3♀). CMV-Cre line: wild-type (n= 13♂ and 13♀) or gHT (n= 8♂ and 5♀). Statistical analyses for B-D are via One-way ANOVA with Tukey’s post-hoc test. Statistical analyses for E-G are via unpaired Student’s t-test. ^*^ p < 0.05; ^**^ p <0.01.

To examine learning and memory using a different testing paradigm, we performed rotarod testing, which primarily assesses motor function, but can be used to address motor learning. Mice are placed upon a rotating rod that incrementally increases in speed (Fig. S4A). Over the course of repeated testing, mice learn to avoid falling from the rotarod. *Chd4* cHT and cKO mice did not exhibit any differences versus controls, nor did we observe any difference in male versus female performance (Fig. S4B, C). Likewise, we did not observe any defect in *Chd4* gHT animals in the rotarod test when sexes were pooled (Fig. S4D). However, when sexes were separated, male gHT mice did not show any phenotype on the rotarod, but female gHTs exhibited impaired motor learning over the course of rotarod training (Fig. S4E). Since *Chd4* cHT animals do not exhibit corresponding phenotypes, these data indicate that female gHTs exhibit learning deficits that map to neurodevelopmental events that map outside of the *Emx1-Cre* expression window.

## Discussion

NDDs are highly heterogeneous – both symptomatically and in terms of etiology. At the genetic level, chromatin remodellers constitute a large proportion of the risk genes linked to these disorders, but the molecular mechanisms that link genotype to phenotype are not well understood. Despite a number of landmark studies examining *CHD4* in neurodevelopment, multiple gaps in our understanding of SIHIWES were unresolved. First, while *CHD4* mutations are clearly linked to both behavioral abnormalities and altered cortical growth, it remained unclear to what degree these phenotypes were linked. Indeed, while Chd4 protein is prominently expressed in dividing cortical progenitor cells, it is also expressed in postmitotic neurons and glia (24). Second, while the observed macrocephalic and microcephalic phenotypes implicate *CHD4* in neocortical development, behavioral abnormalities could also map to other brain regions. Indeed, previous work in the murine cerebellum has already demonstrated that *Chd4* is required for behavior independently of effects on brain growth (14, 23). Third, an important caveat is that previously reported *Chd4* mutant phenotypes manifested in cKOs rather than in heterozygotes. Since human *CHD4* mutations are exclusively heterozygous (16–18), it remains unclear to what degree these phenotypes actually reflect SIHIWES. A fourth related question is whether SIHIWES mutations are loss-of-function alleles, or pathological gain-of-function variants.

To begin to address these research gaps, we generated novel mouse models. Ablation of *Chd4* specifically in the developing telencephalon using the *Emx1-Cre* driver permitted survival of cKOs into adulthood, enabling the cortical knockout phenotype to be assessed postnatally for the first time. We predicted that germline versus conditional mutants would present with similar neurodevelopmental deficits that would differ only in severity. Instead, the observed phenotypes were categorically different – at both the neuroanatomical and behavioral levels. These results illustrate the pleiotropy of chromatin remodelling genes in neurodevelopment, and underscore the requirement for carefully matching genetic models to human NDD etiology.

### Chd4 loss-of-function impairs cortical growth

Using the *Nestin-Cre* driver to generate *Chd4* cKOs, Nitarska et al. previously characterized defects in neocortical progenitor proliferation and apoptosis, culminating in perinatally lethality (24). We obtained the same conditional *Chd4* allele used by Nitarska et al. (generated by the Katia Georgopoulos lab (1)), but instead utilized the *Emx1-Cre* driver, which is expressed in a later and more restricted fashion. We found that *Chd4* cKOs were viable into adulthood, albeit at reduced Mendelian frequencies. cKOs exhibited a prominent reduction in cortical size that was similar to that reported by the Riccio lab (24). The observed microcephalic cKO phenotype also resembles growth impairments reported in other NuRD cKOs, including *Mbd3* or *Hdac1/2* (27, 28). Biochemical data additionally suggest that Chd4 is particularly required in the context of the NuRD complex in embryonic cortical progenitor cells, when Chd3 and Chd5 are less expressed (24).

Surprisingly, despite the fact that cHTs and gHTs are both hemizygous for *Chd4* throughout neocortical neurogenesis, gHTs exhibited microcephalic phenotypes while cHTs were normal, except for a relatively subtle expansion in white-matter and oligodendroglial cells. The lack of analogous growth phenotypes in cHTs suggests that the requirement for *Chd4* during the neurogenetic developmental window is resilient to haploinsufficiency. Instead, the microcephalic phenotype in gHTs likely arises due to cortex-extrinsic pathways, or from earlier neurodevelopmental functions. *Emx1-Cre* -mediated gene excision begins at approximately E10.5 (25), bypassing early telencephalic development. Indeed, in addition to its role in NuRD, Chd4 may also contribute to neurodevelopment through the ChAHP complex (10). ChAHP is implicated in early neurodevelopment, since *Adnp* knockout mice were reported to die at approximately E8.5 and exhibited defective neural induction and gene expression in the forebrain (29). Analogous effects were seen in pluripotent stem cell lines (10, 30, 31). Moreover, *ADNP* is one of the most frequently mutated genes associated with ASD/ID (32, 33). However, ASD has only been linked to one SIHIWES individual so far (17). Future work will be required in order to clarify the contribution of NuRD and ChAHP chromatin remodelling complexes to neurodevelopment.

Although some SIHIWES patients have been reported to exhibit microcephaly (17), macrocephaly has been more frequently observed. Along with the Nitarska et al. study (24), our observations utilizing strong loss-of-function mutants argue that macrocephalic *CHD4* alleles likely act via gain-of-function effects rather than via hemizygosity. Most SIHIWES alleles described to-date presented with *de novo* missense pathogenic variants – usually located within the ATPase/helicase domains (16–18). These variants might generate enzymatically-inactive proteins that might nonetheless incorporate into chromatin remodelling complexes and bind regulatory elements in a non-productive manner.

In terms of neocortical cell composition, *Chd4* cKOs had a significant reduction in Brn2+ and Satb2+ upper-layer cortical neurons, similarly to the perinatal phenotypes reported by Nitarska et al. The more prominent reductions in late-born upper layer neurons might reflect the requirement for chromatin remodelling during a specific temporal window of neurogenesis. Alternatively, growth defects might be cumulative over time, such that they have a disproportionate effect at late stages.

### Germline versus conditional mutations lead to divergent effects on behavior

We performed extensive behavioral testing in order to functionally validate our models. With respect to cognition, SIHIWES mutations are heterogeneous. For example, a pair of individuals harboring the same Arg1127Gln mutation had markedly different cognitive abilities. One individual exhibited moderate ID, and the other had IQ scores within the normal range (17). Perhaps accordingly, we found that each *Chd4* mouse model presented with a different subset of behavioral defects, which are likely dependent on where and when *Chd4* expression is altered. Changes in repetitive behaviors and anxiety were affected only when the cortex was conditionally targeted, while changes in learning behaviors occurred only when the whole body was targeted – and only in female mice.

The connection between behavioral phenotypes and biological sex was unexpected. It is unclear why females would be more affected, though sex-specific changes in learning behaviors have previously been reported in other rodent NDD models (34–37). Female-specific phenotypes were mainly observed in gHT mice, and correlated with microcephaly. However, marble-burying phenotypes in cHT and cKO animals also appeared to track with sex, further hinting at increased vulnerability to chromatin remodelling dysfunction in females. In humans, SIHIWES cases only include 12 females and 20 males, making it hard to draw conclusions regarding sex-specific differences. Males are generally diagnosed with NDDs more frequently than females, although this is partly attributable to the importance of X-linked genes within the genetic landscape of these disorders.

We tested animals for behavioral deficits that might be relevant to NDDs, but also examined animals for activity levels, sensory deficits, and motor system deficits. cHT, cKO, and gHT mice exhibited intact optomotor reflexes, indicating normal visual function. All mutants exhibited equivalent circadian activity patterns that were not different versus controls. cKOs had slightly reduced body weight versus control animals, but we did not detect other obvious health defects in these animals, except that a subset of cKO adults developed skin lesions on their paws, which we believe may be due to *Emx1-Cre* expression in the limbs (25, 38). We also observed hydrocephalus in at least one *Chd4* cKO animal. While hydrocephalus is linked to SIHIWES, the frequency of this observation in cKO mice was too low to determine whether it was truly linked to genotype. We excluded hydrocephalic animals from our behavioral testing.

While cKO animals had the strongest anatomical and behavioral deficits, they likely fail to capture certain relevant phenotypes – such as deficits in spatial and motor learning. Despite the fact that gHT animals have the most ‘construct validity’ in terms of their resemblance to SIHIWES mutations, behavioral deficits in these animals were relatively mild, and were confined only to female mice. Moreover, macrocephaly was not observed. Again, these data argue that SIHIWES mutations may not represent simple loss-of-function mutations. In the future, it will be important to generate mutants harboring patient-specific mutations, in order to shed more light on pathogenic mechanisms and SIHIWES etiology.

## Methods

### Animal Work

All animal work was approved by the University of Ottawa Animal Care Committee and carried out following guidelines set out by the Canadian Council of Animal Care. The University of Ottawa Animal Care and Veterinary Services facility provided animal housing, where animals were kept in a 12-hour light cycle with food and water provided *ad libitum*. *CMV-Cre* mice (26) were purchased directly from the Jackson Laboratory (B6.C-Tg(CMV-cre)1Cgn/J, Stock no. 006054). *Emx1-Cre+* mice (25) were generously donated by the David Picketts laboratory (OHRI, Ottawa, Canada). *Chd4^Flox/Flox^* founders were generously donated by the Katia Georgopoulos laboratory (Harvard). *Emx1-Cre* mice and *Chd4^Flox/Flox^* mice were backcrossed to C57BL/6J background mice a minimum of six generations before experimentation began. Animals were genotyped using primer sets published in the referenced papers.

### Immunofluorescence

Adult mice were euthanized via CO2 inhalation followed by cervical dislocation. The brains were dissected out, weighed, imaged, and fixed in 4% PFA overnight. Mice pups were euthanized via decapitation, followed by brain dissection, imaging, fixing in 4% PFA overnight. Brains were subsequently transferred to 20% sucrose for 24hours, freezing solution (1:1 of OCT:30% sucrose in PBS) for 24-36hours, and frozen down to −80°C in freezing solution.

Brains were sectioned coronally using a Leica CM1860 cryostat. Thickness was set at 14μm for pre-natal brains and 16μm for post-natal brains. Sections were rinsed in 1X PBS followed by antigen retrieval in citrate buffer (10 mM Sodium citrate, 0.05% Tween 20, pH 6.0) using a pressure cooker for 10min, and rinsing with water for 10 min. All antibodies were diluted in 30% w/v Bovine Serum Albumin, 0.4% Triton, and 1:5000 Hoechst in PBS. Primary antibodies were: rabbit anti-Aldh1l1 (1:100, Abcam ab87117), Rabbit anti-Brn2 (1:200, Cell Signaling Technology 12137S), mouse anti-Chd4 (1:200, Abcam ab70469), rabbit anti-Chd4 (1:100, Abcam ab72418), rat anti-Chd4 (1:100, BioLegend 942302), rat anti-Ctip2 (1:200, Abcam ab18465), rabbit anti-Neurog2 (1:250, Cell Signaling Technology 13144S), Rabbit anti-Olig2 (1:200, Novus Biologicals NBP1-28667SS), mouse anti-Pax6 (1:10, Developmental Studies Hybridoma Bank PAX6), Rabbit anti-Pax6 (1:500; Novus NBP2-19711), rat anti-Satb2 (1:100, Abcam ab51502), goat anti-Sox2 (1:200, R&D Systems AF2018), Rabbit anti-Tbr1 (1:100, Cell Signaling Technology 49661), and rabbit anti-Tbr2 (1:100, Abcam ab23345). Primary antibodies were incubated overnight at 4°C, followed by three 1X PBS washes, secondaries (Alexa Fluor 488 goat anti-mouse IgG (Jackson ImmunoResearch 115-545-003; 1:1000), DyLight 550 donkey anti-rat IgG (Invitrogen SA510027; 1:1000), Dylight 650 donkey anti-rabbit IgG (Novus Biologicals NBP1-75634; 1:1000)) for 1 hour at room temperature, three 1X PBS washes and mounted in Mowiol mounting media (12% w/v Mowiol 4-88, 30% w/v glycerol, 120mM Tris-Cl pH 8.5, 2.5% DABCO).

### Cell Counting and Imaging

For consistency across coronal sections, the primary somatosensory cortex located between the crossing of the corpus collosum to the beginning of the hippocampus was targeted. Every other section, barring imperfections, was imaged using the Zeiss LSM900 Confocal Microscope at 200X magnification. Single Z-stack images were tiled and stitched using Zen software (Zeiss). Manual cell counting was performed using Adobe Photoshop on 200μm-wide sections of somatosensory cortex between the crossing of the corpus callosum and the start of the hippocampus. Each 200μm-wide section was separated into six 200μm-wide bins of equal height for counting. This allowed the proportion of cells in the upper sixth area of the cortex to the lower sixth area of the cortex to be assessed, as well as total (i.e. absolute) counts. Three technical replicates were counted for each biological replicate (animal). Images were additionally processed using Photoshop, Powerpoint (Microsoft), and Biorender (Biorender.com).

### Cortex and Cerebellum Area Measurements

Brains were imaged post-dissection using a Zeiss Stemi 508 stereo microscope with Axiocam ERc 5s at 6.3X magnification. Images were imported into Fiji and the outline of cortex or cerebellum was traced using the “polygon selections” tool. Area measurements were recorded and used for subsequent analysis. Scale was consistent across images and is depicted with arbitrary units.

### Statistics

Data is presented as mean ± standard deviation unless otherwise indicated. Round black circles represent individual data points. *Emx1-Cre* vs *CMV-Cre* lines were assessed separately due to significant differences between the control groups of both lines. Statistical analysis was performed vs Student’s t-test, or one-way ANOVA with Tukey’s or Bonferroni post-hoc tests, as indicated in the text. P < 0.05 was considered statistically significant.

### Behavior Analysis

Behaviour testing, including protocols and materials provided, was performed through the Behavioural Core Facility at the University of Ottawa. All testing was blinded to animal genotypes. Mice were assessed between 3-6 months of age, with females and males housed separately. Mice were habituated to the testing room for at least 30 minutes prior to testing commencing. Mice were handled (lifted out of cage and put back 3-4 times) for 5 days prior to commencing. Testing was scheduled so that mice only performed one behavior test a day following established protocols. Behavior testing was recorded with a video camera mounted appropriately and subsequently analyzed using Ethovision software (Noldus). Velocity and distanced travelled measurements were captured during each behavior test. During analysis, male and female data was pooled, unless otherwise indicated.

#### Elevated Plus Maze

Mice were placed in the middle of a plus shaped apparatus that was elevated above the ground for 10 minutes under 100 lux lighting. Two arms of the plus maze, opposite to each other, were enclosed (6 cm wide × ~69cm long, 20 cm high walls) while the remaining two were open (6 cm wide × ~69cm long, raised 74 cm off the floor). The number of entries the mouse made into the closed versus open arms, as well as the time spent in the closed versus open arms was measured. Measurements of mice activity, videos of mice activity, and analysis was collected using Ethovision software (Noldus).

#### Open Field

Mice were placed in the middle of an opaque plastic box (45 cm × 45 cm × 45 cm) with an open top. A camera mounted above the box recorded the mice activity for 10 minutes under 300 lux light. Ethovision software (Noldus) was used to analyze videos of mice activity. The amount of time a mice spent in the center of the box (small centre and large centre) and the corners of the box was recorded. The latency for the mice to reach the corners of the box was also measured.

#### Beam Break

Mice were placed in a new housing cage that was placed between a photocell emitters and photocell receptors system for 24 hours. Disruption of the infrared light as the mouse moved through the cage was detected and recorded by the photocell analyzer (Micromax Analyzer, AccuScan) and subsequently tallied through computer analysis (Fusion 5.3 software, Omnitech Electronics Inc.). Ambulatory Activity Counts, measuring the amount of beam breaks that occurred while the animal was ambulating (vs stereotypic behaviour such as grooming, etc.) was plotted.

#### Marble Burying

Mice were first habituated to the testing arena with no marbles present for 5 minutes. Subsequently, 20 marbles were evenly spaced atop woodchip bedding in a 4 × 5 marble grid in the arena. Mice were placed in the testing area again for 30 minutes and the number of marbles buried after this time was manually recorded. 75%+ of marbles covered by bedding was considered as buried.

#### Adult Social Interaction

Mice were habituated in red light before testing, and the test was subsequently performed under red light. During the habituation phase, a mouse was placed in the corner of a 45 cm × 45 cm × 45 cm box with a 5.5 cm × 9.6 cm wire mesh rectangular cage located along one side and allowed to explore for 5 minutes. After this point, the mouse was removed and temporarily placed in a clean container for 5 minutes. Next, a C57BL/6J background social target mouse (of the same gender and of similar age) was placed inside the wire mesh cage for the socialization phase. The tested mouse was reintroduced into the box for 5 minutes and activity recorded. Time spent and number of entries of the mouse’s body near the cage (large interaction zone) and in the corners was recorded, as well as time and number of entries the nose point of the mouse entered the small interaction zone around the cage.

#### Rotarod

Mice were placed on a horizontal textured rod (IITC Life Science, Ugo Basile) rotating at a speed of 5 RPM. The speed was gradually increased to 40 RPM over the course of five minutes and the time the mouse spent on the rod before falling off (latency to fall) was recorded. Mice were trained on the Rotarod four times a day with a 10 minute intertrial interval in their home cage. This was performed for four consecutive days.

#### Morris Water Maze

A ~132cm diameter pooled was filled with opaque (white non-toxic tempura paint) water (22°C) with a 10cm diameter white platform hidden 1cm below the surface of the water. The pool had 4 starting location at the edge of the pool located equidistant to each other, and the platform was located between two of them about 24cm from the edge of the pool. Lighting was set to 140 lux and the pool was surrounded by 4 white “walls”. Two black visual cues were located on two of the adjacent walls to allow for spatial navigation. Briefly, a mouse was placed in the water at one of the starting points and allowed to try and find the platform for 1 minute. If they did not find it, the platform was tapped by the researcher, and if the mice still did not find it, gently guided by the tail towards the platform. Once on the platform, the mouse was left for 10 seconds, prior to being removed, toweled off and returned to its home cage for a 15-minute intertrial interval. Mice were trained 4 times a day at each of the locations, the order of which was randomized. After 6 days of training, the platform was removed and the amount of time the mouse spent in the quadrant where the platform was located, and the number of times the mouse crossed the area where the platform had been was measured.

#### Optomotor Analysis

The StriaTech Optodrum was used to assess visual acuity in mice. Briefly, a mouse was placed atop a white 9 cm diameter platform in the middle of the OptoDrum (box shaped). Contrast cycles was set at 99.72% with a constant rotation speed of 12°/s. The number of cycles was set to start at 60 cycles (0.167 cyc/°) and increased/decreased following an automatic staircase procedure that allowed determination of visual acuity (i.e., max number of cycles mice would respond to). Mice were tested three times in one day and the mean of these trials was calculated and graphed.

## Author Contributions

Conceptualization: S.L., P.M. Data curation: S.L., P.M. Formal analysis: all authors. Investigation: all authors. Project administration, resources, supervision: P.M. Visualization: S.L., P.M. Writing—original draft: S.L., P.M. Writing, review, and editing: all authors.

## Acknowledgements

We are indebted to Katia Georgopoulos for previously sharing *Chd4^Flox^* mice, and to David Picketts for sharing *Emx1-cre* mice. We are grateful to Karin Weiss for providing helpful advice. We are grateful to Kerstin Ure, Sarah Kealey, and Katherine Tacay from the *Animal Behaviour Core*, for their help with behavior testing. We also thank *Animal Care and Veterinary Service*, and *Cell Biology and Image Acquisition* core facilities. We thank Samuel Clémot-Dupont, Ida Malloy, and Mattar lab members for their ongoing help and input. This project was supported by the Canadian Institutes of Health Research (CIHR) Operating Grants PJT-166032 and PJT-166074. S.L. was supported by graduate scholarships from the Natural Sciences and Engineering Research Council of Canada (NSERC), as well as the Ontario Graduate Scholarship. P.M. currently holds the Gladys and Lorna J. Wood Chair for Research in Vision.

---

**Figure S1.**
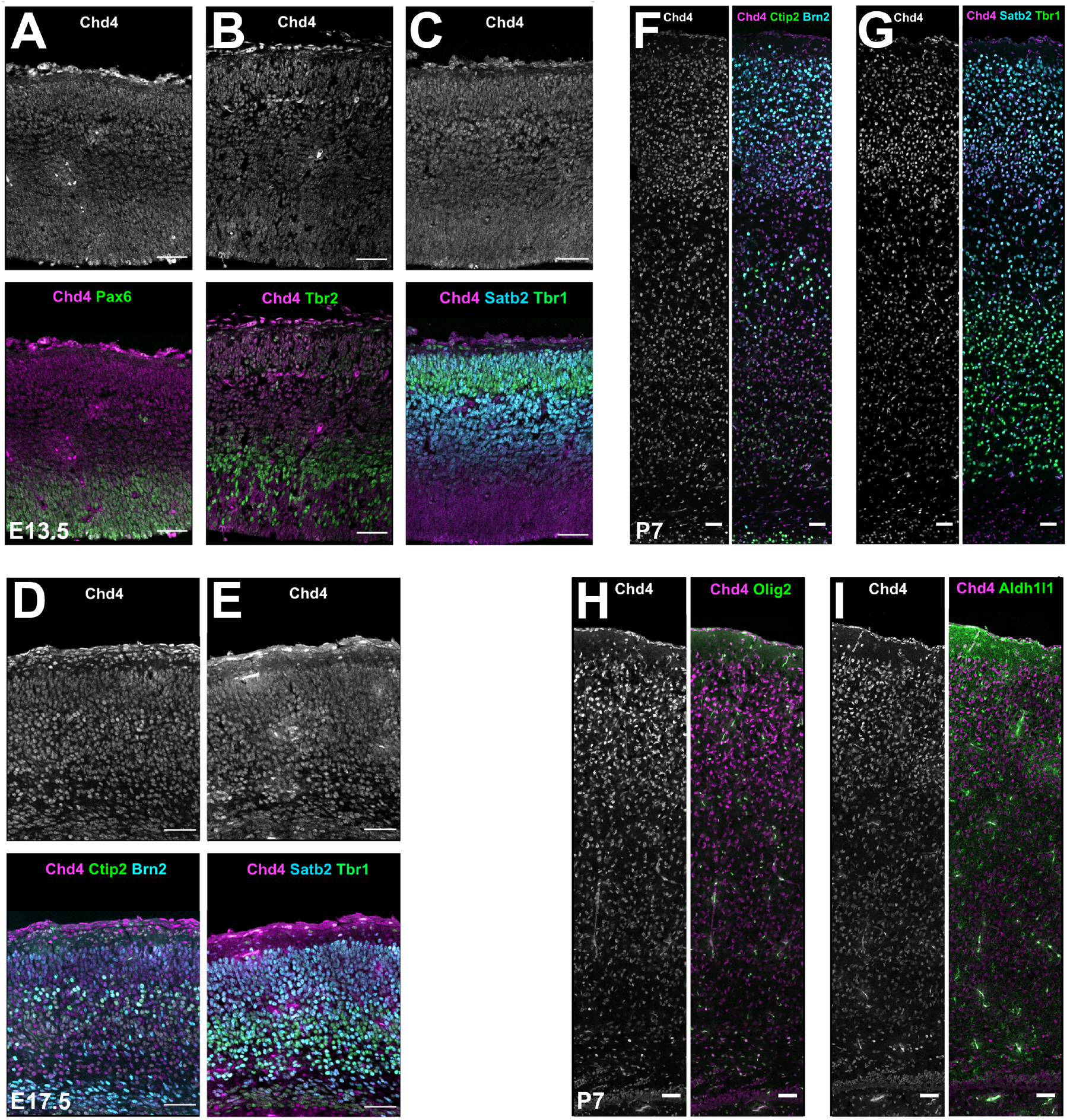
Chd4 expression during neocortical development. (A-H) Immunohistochemistry for Chd4 and cell-type-specific markers as indicated, at E13.5 (A-C), E17.5 (D, E), and P7 (F-I). Scale bars = 50 μm.

**Figure S2.**
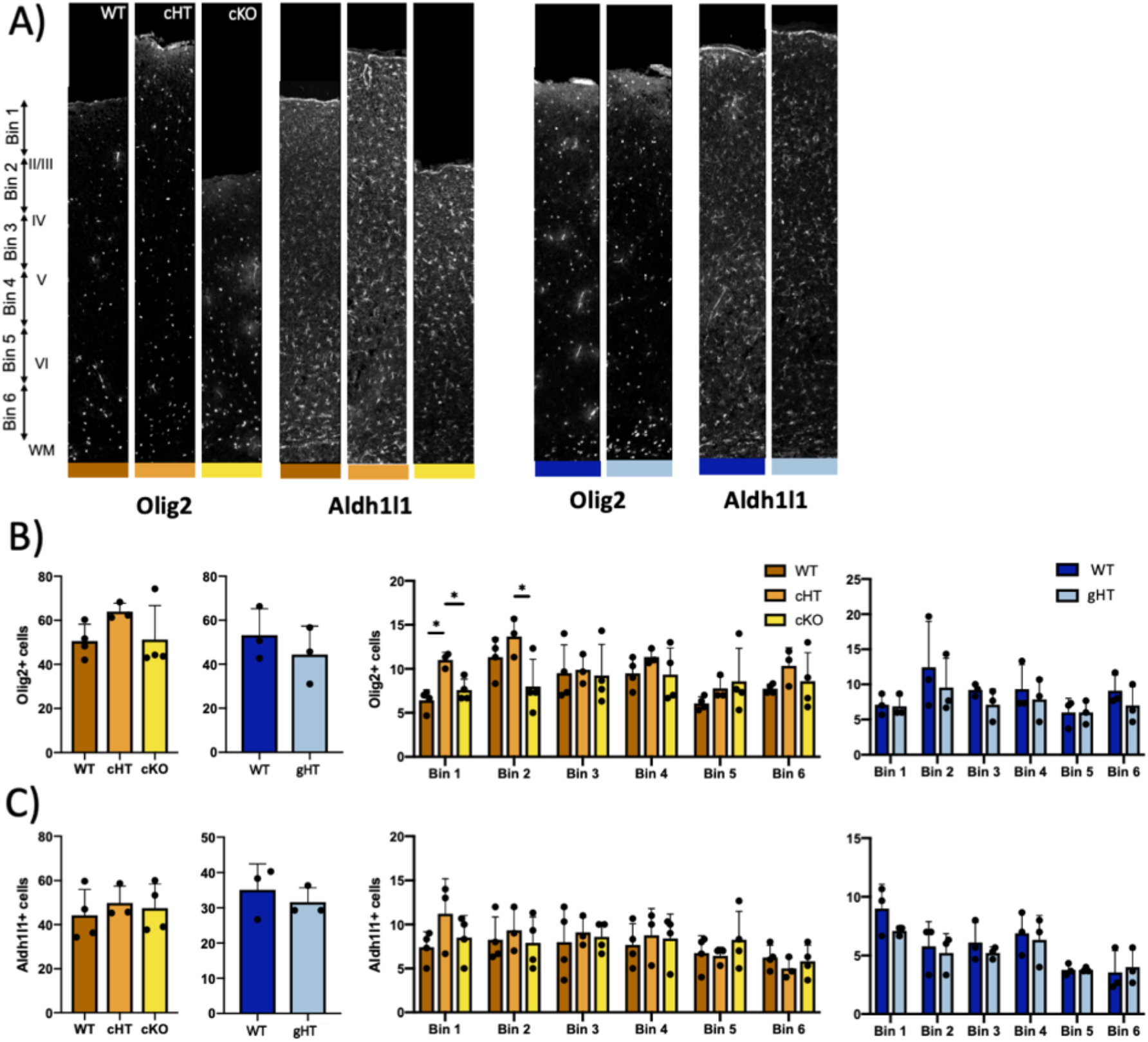
Assessing glial cells in *Chd4* mutants. (A) Immunohistochemistry on coronal sections from wild-type, cHTs, cKOs, or gHTs for the oligodendrocyte-lineage marker Olig2, or the astrocyte marker Aldh1l1. (B, C) Absolute cell counts, or counts taken from 6 equally sized bins (see A) for the indicated genotypes. (B) Olig2. (C) Aldh1l1. Statistical analyses for WT, cHT, and cKO comparisons are via one-way ANOVA with Tukey’s post-hoc test. Comparisons of control vs. gHT are via Student’s t test. ^*^ p < 0.05. ^**^ p < 0.01. *Emx1-Cre* line: WT n=4, cHT n=3, cKO n=4. *CMV-Cre* line: WT n=3; gHT n=3.

**Figure S3.**
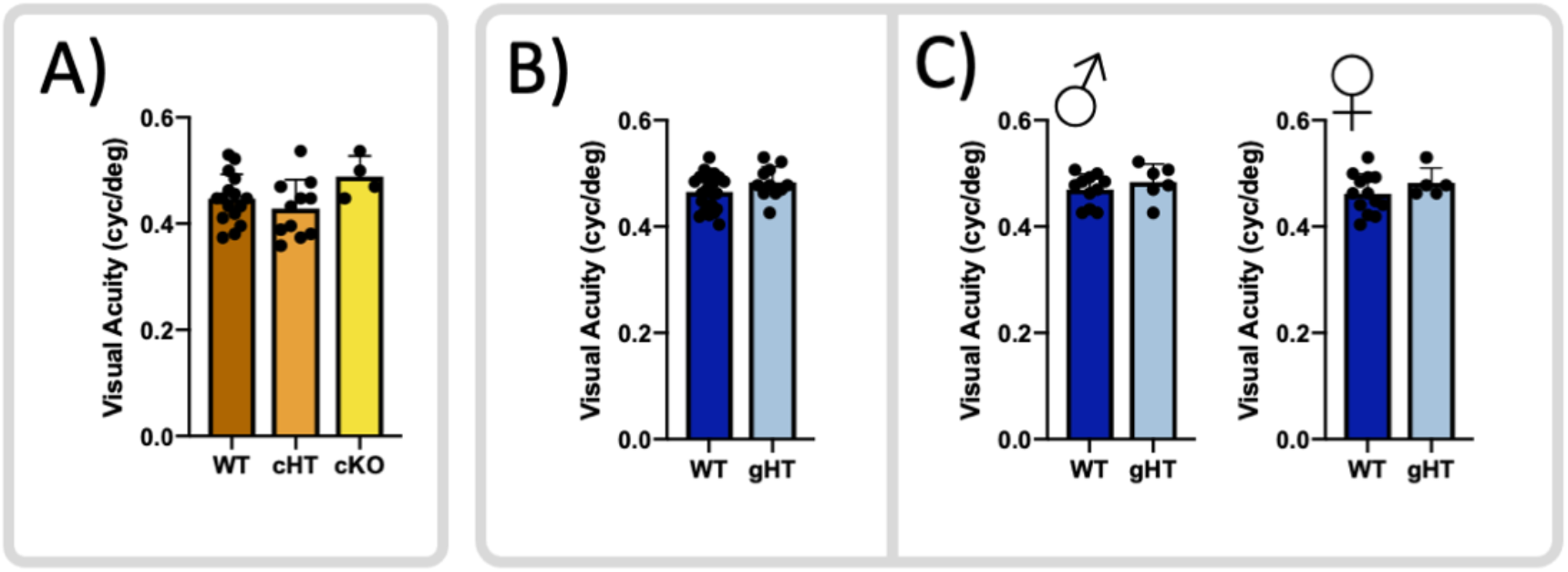
Visual acuity is unchanged in *Chd4* mutants. (A) Visual acuity of WT, *Chd4* cHT and *Chd4* cKO mice. (B) Visual acuity of WT and *Chd4* gHT mice. (C) Same data as in (B) but separated by biological sex. Statistical analyses for WT, cHT, and cKO comparisons are via one-way ANOVA with Tukey’s post-hoc test. Comparisons of control vs. gHT are via Student’s t test. *Emx1-Cre* line: WT n = 16, cHT n=11, cKO n=4. *CMV-Cre* line: WT 11♂ and 13♀, gHT, 6♂ and 5♀.

**Figure S4.**
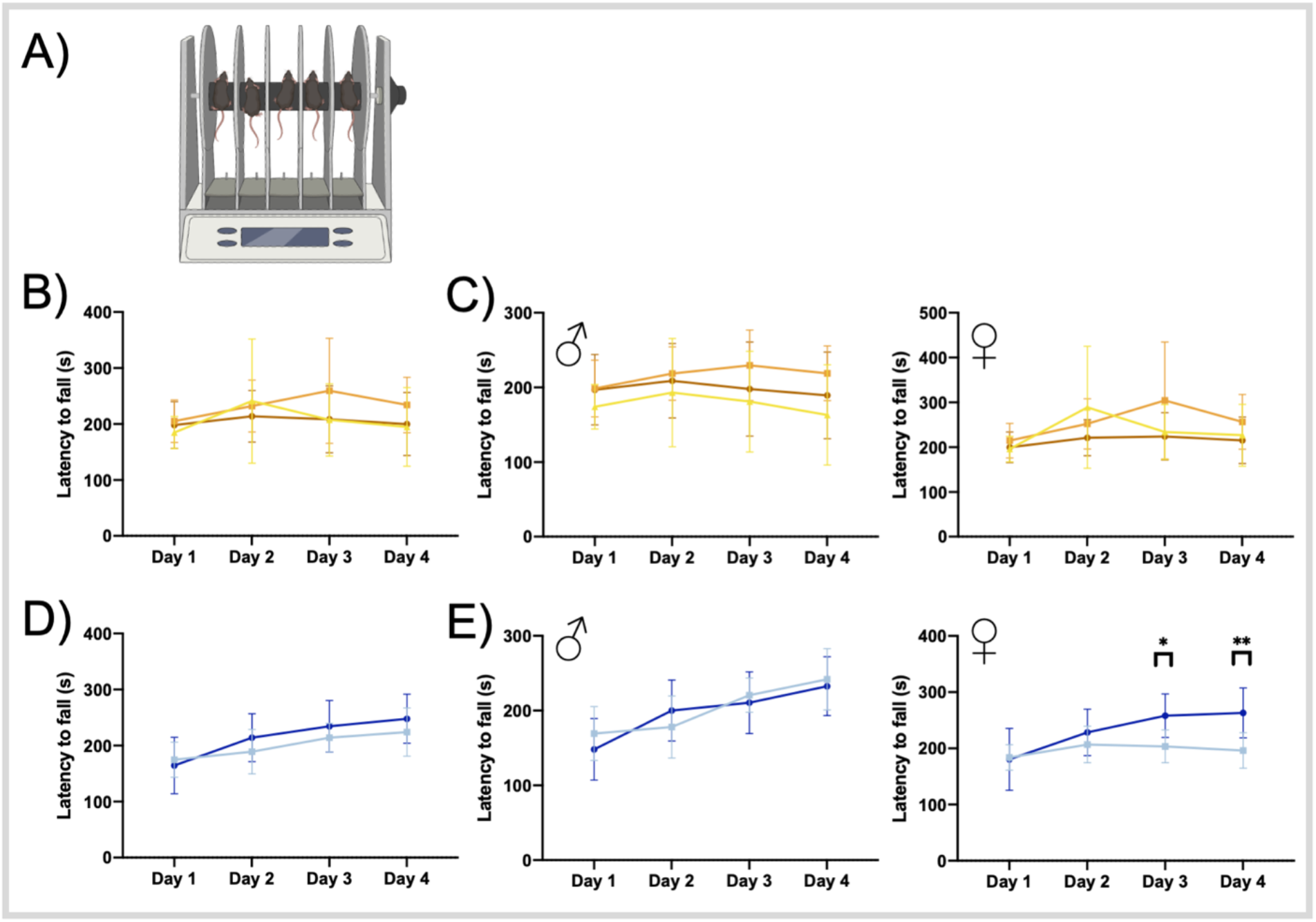
Motor learning behaviors in *Chd4* mutants. (A) Schematic of the accelerating rotarod apparatus. Testing was performed for 4 consecutive days with 4 trials per day, 5 minutes each. (B) Latency to fall for wild-type (n= 19♂ and 13♀), cHT (n= 9♂ and 6♀), and cKO (n= 3♂ and 3♀). (C) Same data separated by sex. (D) Latency of fall for wild-type (n= 13♂ and 13♀) or gHT (n= 8♂ and 5♀). (E) Same data separated by sex. Statistical analyses for WT, cHT, and cKO comparisons are via one-way ANOVA with Tukey’s post-hoc test. Comparisons of control vs. gHT are via Student’s t test. ^*^ p < 0.05. ^**^ p < 0.01.

